# Theta-Gamma Phase-Amplitude Coupling Supports Working Memory Performance in the Human Hippocampus

**DOI:** 10.1101/2024.03.24.586454

**Authors:** Yousef Salimpour, William S. Anderson

**Affiliations:** Functional Neurosurgery Laboratory, Department of Neurosurgery, Johns Hopkins School of Medicine, Baltimore, Maryland, USA

**Keywords:** Cross-frequency coupling, phase-amplitude coupling, electrocorticography, working memory, sequence learning, human memory structures, epilepsy disorder

## Abstract

Phase-amplitude coupling (PAC) occurs in the human hippocampus during working memory and supports the contribution of the hippocampus in the maintenance of multiple items. Additionally, PAC has the potential to reveal the neural mechanisms underlying multi-item maintenance in the hippocampus by providing a putative architecture for multi-item representation. Theta and gamma range rhythms are prominent neuronal oscillations in the hippocampus. Studies on the role of theta frequency oscillation in local field potentials in human memory have shown mixed evidence for successful remembering. The role of gamma oscillatory activity in contributing to memory retrieval is not yet fully understood. They also interact with each other in the form of PAC during memory performance. This study aims to investigate the neurophysiological function of theta-gamma PAC in the human hippocampus during a multi-item working memory task and characterize its association with performance. Theta-gamma cross-coupling investigation in the electrocorticographic signals was performed from the hippocampus recording of ten epilepsy patients while they were engaged with the working memory task. The results show strong correlations between PAC levels and the subjects memory performance, but no correlation with theta and gamma power individually, specifically in the retrieval phase of a working memory task. These observations demonstrate the possible role of PAC in memory-related operations, suggesting a PAC-based neural mechanism for working memory in the hippocampus.

**Significance Statement:** The findings from this study elucidate the crucial role of phase-amplitude coupling in the human hippocampus during working memory tasks, specifically in the maintenance of multiple items. By analyzing electrocorticographic recordings from epilepsy patients engaged in a working memory task, our research unveils a direct correlation between PAC levels and memory performance during the retrieval phase, which is not observed when analyzing theta and gamma oscillations individually. These findings suggest a theta-gamma coupling based mechanism within the hippocampus that facilitates working memory, offering new insights into the complex neural processes underlying memory encoding and retrieval. This advancement in understanding the neural architecture of memory not only contributes to the foundational knowledge of cognitive neuroscience but also opens avenues for developing targeted interventions for enhancing memory performance with translational application in treating memory-related neurological disorders.

## 1 Introduction

The human hippocampus is a prominent cortical structure in the medial temporal lobe known to be essential for working memory function. Neuronal oscillations are observed across various spatial and temporal scales in the hippocampus (Canolty et al., 2010) in healthy subjects and patients with neurological disorders (Salimpour and Anderson, 2019; Hyafil et al., 2015; Goutagny et al., 2013; Wang, 2010). During the increased memory-related activity, theta and gamma rhythms are generated by neural circuits in the hippocampus. These neural rhythms are a neuronal mechanism for organizing hippocampal spike timing using different temporal scales in the encoding and retrieval of information during memory tasks (Roux et al., 2014). Specifically, invasive electrophysiological recordings in the human hippocampus demonstrated persistent neuronal activities during working memory maintenance in the single cell (Kaminski et al., 2017), cortical oscillations (Brzezicka et al., 2019), and cross-frequency coupling (Axmacher et al., 2010)

In the hippocampus, theta band oscillation is generated by a contribution of the local activity appearing from the spontaneous interaction of interneurons and pyramidal cells and different types of input, such as GABAergic, cholinergic, and glutamatergic (Goutagny et al. 2009; Gloveli et al. 2005). One of the suggested roles for theta oscillations in the hippocampus is the separation of neuronal processes associated with the encoding of new information and the retrieval of stored information (Hasselmo et al., 2002). In the normal activity state, the memory-associated structures must be able to organize and navigate the access to the new sensory information and the retrieval of stored information. Theta range oscillations might be a neuronal mechanism to coordinate cell assembly to reduce the likelihood of interference between encoded and retrieved information (Hasselmo et al., 2002; Hasselmo, 2005; Colgin and Moser, 2010). Particularly, the theta phase modulates the inputs to the hippocampus, according to recent animal studies. The new input information (mostly from the entorhinal cortex) is at the highest rate at the trough of theta oscillation in the hippocampus which is associated with the encoding phase of memory processing (Brankack et al., 1993; Kamondi et al., 1998). On the other hand, CA1 neurons are modulated by CA3 inputs at the highest rate indicating more internal process and information retrieval (Hasselmo, 2005). These phase-specific physiological alterations allow the hippocampus networks to control the direction of various types of memory-related information depending on encoding or retrieval function.

The gamma rhythm is another main rhythmic activity of the hippocampus has a limited understanding of its neurophysiological mechanisms. There are pieces of evidence for the contribution of the interneurons and pyramidal cells in gamma oscillations which may serve to balance inhibition and exhibition (Csicsvari et al. 2003; Buzsaki and Wang 2012). During multisite recording studies, when CA1, CA3, and EC are recorded simultaneously, oscillations in the high gamma range are an indication of improved coordination between CA1 and EC representing ongoing neuronal encoding, whereas oscillations in the low gamma range show improved coordination between CA3 and CA1 which might be associated to the information retrieval process. These oscillations also often occur on distinct cycles and at different theta phases (Colgin et al., 2009). Other investigations have found task-dependent hippocampus activity modulation, which is consistent with the retrieval of stored memories (Douchamps et al., 2013; Newman et al., 2013). These results imply that new information is encoded and stored information is retrieved using a complicated neuronal process that involves a complex interaction between neuronal populations.

Additionally, oscillations of different frequencies in the hippocampus demonstrate complicated coupling properties reflecting more complex features of neural oscillations present in memory-related activity, mostly in the form of cross-frequency coupling metrics (CFC) (Roux et al., 2014). One of the earliest illustrations of CFC in the hippocampus was between the amplitude of gamma rhythm and the phase of the theta oscillatory activity, which is called the phase-amplitude coupling (PAC) (Canolty et al., 2006; Tort, et al. 2007). Although the neural mechanism underlying PAC in the hippocampus is not fully understood, current pieces of evidence show the contribution of both pyramidal cells and interneurons, which fire simultaneously at specific phases of theta (Klausberger et al. 2003; Tukker et al.2007)

Investigating the role of theta and gamma CFC in memory formation and retrieval has immense value for both basic science and clinical applications. In combined with existing computational modeling of CFC generation, the putative theta-gamma coupling architecture for multi-item representation has the potential to investigate the underlying neural mechanism for working memory in the hippocampus. Recent evidence from human studies has demonstrated that the theta phase and gamma amplitude CFC in the hippocampus are correlated with short-term memory performance (Axmacher et al.2010; Fell and Axmacher, 2011). To this end, we demonstrate the association between CFC (in PAC format) and working memory performance by analyzing the cortical electrocorticogram (ECoG) signals recorded from the hippocampus of epilepsy patients during a visual memory task. We evaluated the memory performance and measured PAC during the retrieval phase of the memory task. We focused on theta phase and low-gamma oscillation during the retrieval period. These observations may demonstrate the possible contribution of CFC in memory formation and retrieval during normal function of the nervous system and possible correlations with neurological disorders (Fujita et al., 2022; Bazzigaluppi et al., 2018).

## 2 Materials and Methods

A previous study from our laboratory consisted of measuring ECoG signals in the gamma frequency band when a subject was asked to recall a sequence of images over a certain timespan (Madhavan, et al., 2015). This study proved that the power in the gamma range correlated with memory performance during the recall phase (Madhavan, et al., 2015). In the current effort, we used a similar visual memory paradigm, including similar image sets and multiple trials. Still, here we focus on CFC in general and PAC specifically to explore the function of the hippocampus. The cross-coupling between the amplitude of the gamma oscillation and the phase of the theta rhythm is the main outcome measure derived from memory testing.

### Human Subjects

Ten participants with epilepsy (6 males, mean age 40.1 ± 7.4 years) took part in our study. They were implanted with depth electrodes as part of their diagnostic assessment for pharmaco-resistant focal epilepsy and the recordings were performed at Johns Hopkins Hospital. The placement of electrodes was based on clinical needs for localizing the seizure onset zone and considering the feasibility of surgical resections. To be eligible for the study, participants had to have normal IQ, and electrodes implanted in the hippocampus contralateral to or outside of the epileptogenic region were included for the analysis of recorded ECoG signals. All steps of the research project, including the behavioral task and data collection procedures, were approved by The Institutional Review Board at the Johns Hopkins University School of Medicine. Written informed consent was obtained from all participants before behavioral testing and electrophysiological data collection. The clinical characteristics of all subjects in this study are summarized in Table I.

### Anatomical localization and channel selection

Based on the subjects’ neuroimaging data, an independent neurologist visually determined all electrode locations. We recreated the electrode placements in the interested target to provide further visualization. Implanted electrodes were anatomically localized using the pre-implant T1-weighted MRI and a post-implant CT scan. The Brainnetome Atlas was used for anatomical labeling of the regions of interest (Fan et al., 2016). The post-implant CT scan was co-registered with the MRI using the FieldTrip toolbox (Griffin et al., 2018). Electrodes that are visible in the CT were localized to specific structures manually and verified by a neurologist with expertise in anatomical structures. The electrode coordinates were transferred to the MRI coordinates and mapped to the standard space to enable group visualization. Successful anatomical localization provided anatomical labeling for individual contacts and facilitated the visualization of the electrode coverage in all subjects via a standardized brain. One pair of hippocampal depth electrode contacts was selected per participant from each side of the brain based on anatomical and functional criteria. The selection of channels was done in individual subject coordinates to avoid errors due to distortions while converting in MNI space. Channels of interest were selected only if they were located in the hippocampus, and in the case of bilateral hippocampal recordings, only one pair of channels was chosen for each side. Channels were referenced to the adjacent more distal contact along the electrode using bipolar referencing. To visualize the selected contacts across our sample, we normalized each participant’s pre-operation MRI to MNI space.

Finally, to facilitate the visualization of contacts across the group, a sphere was placed around each contact’s center point and overlaid across the participants (Fan et al., 2016; Griffin et al., 2018).

### ECoG recording protocol

The clinical research team implanted epilepsy subjects with multiple intracranial electrodes, including grids and strips (Ad-Tech Medical, Racine, Wisconsin, and PMT, Chanhassen, MN), directly on the surface of the cortex. Additionally, in most of the study subjects, multi-contact cylindrical depth electrodes with platinum contacts were implanted for diagnostic purposes as well (Ad-Tech Medical, Racine, Wisconsin). Ictal localization was performed to guide a possible subsequent surgical resection after identifying the spatial seizure onset zone. After undergoing the neurosurgical procedure for implantation of the electrodes, patients underwent one to two weeks of electrocorticography monitoring in the epilepsy monitoring unit with additional concurrent video observation.

### Visual Memory Task

For measuring the memory performance of our subjects, we used our previously designed visual memory task (Salimpour and Anderson, 2019; Madhavan, et al., 2015). In brief, all our subjects took part in a four-image version of an image sequence recall task. They performed the image sequence recall task implemented in Psychophysics Toolbox version 3 (Kleiner et al., 2007; Brainard, 1997) using the MATLAB platform. Prior to each recording section, participants were informed that an image sequence consisting of four images would be presented and after that, the ordering of the displayed images would be later requested (encoding phase: about 6s). After a delay period (1500ms), their memory performance was tested for a portion of the sequence order (two images each time) two times at each trial (retrieval phase: about 6s) (Figure 1). Subjects were asked to point to the image they believed had appeared earlier in the sequence using one of two assigned buttons on a gamepad controller. During each testing session, if within a minimum of 20 trials, the subject achieved eight consecutively correct recalls, the stop criterion was met, and the testing was terminated. Otherwise, they continued with repeated trials until they were able to meet the criteria or were stopped by the experimenter. Each session included a brief pre-training phase to ensure that participants memorized the sequence. All the task timings were recorded and tagged using a serial port. In each trial, we recorded and analyzed ECoG data in three phases of the task, including the baseline phase, encoding phase, and retrieval (recall) phase (see Madhavan et al., 2015 for more detail). However, in this study, our focus was mostly on two recall phases (Figure 1.B).

**Figure 1.**
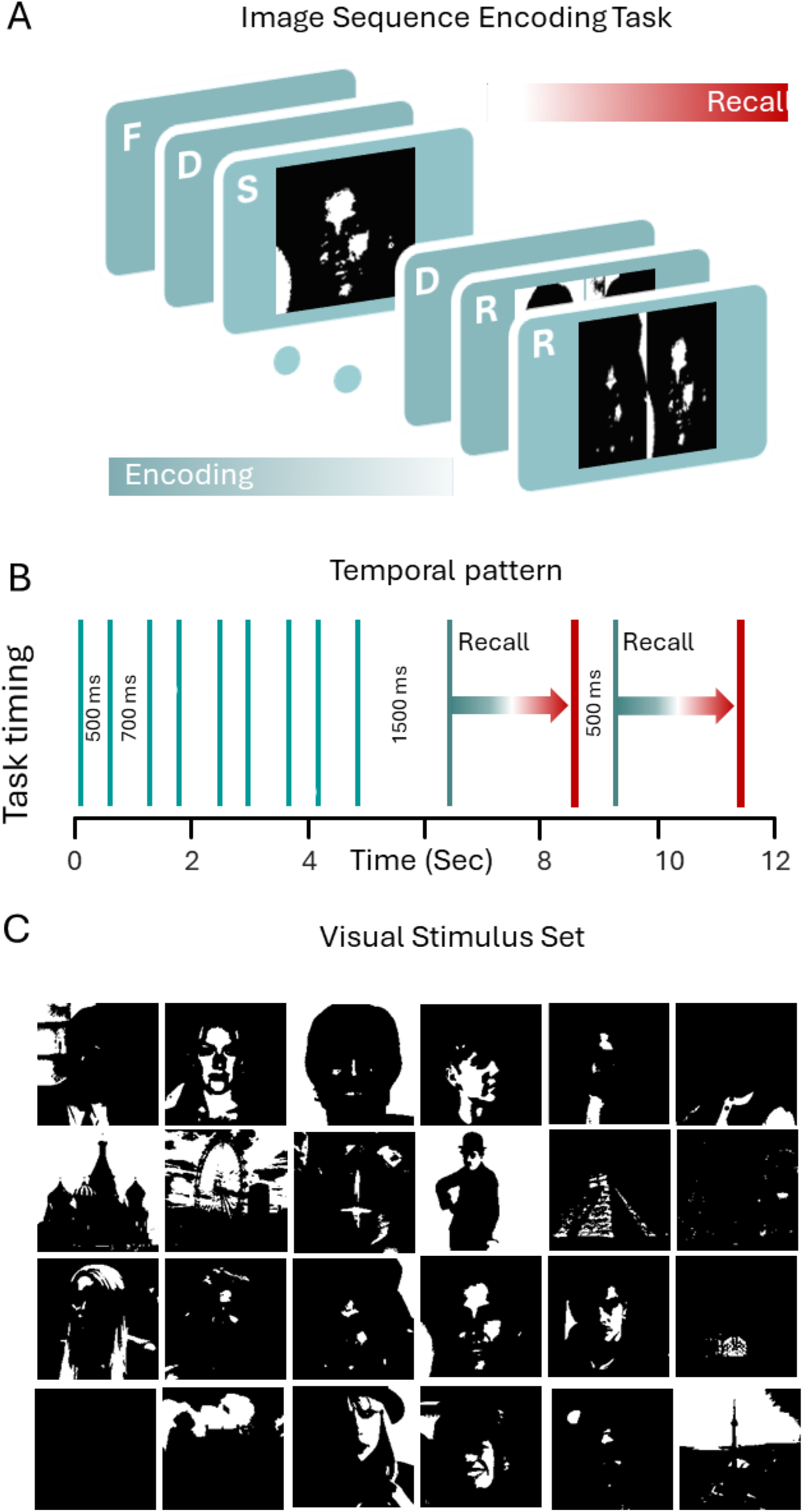
Visual memory task. **A.** The image sequence recall paradigm involves a trial that begins with a fixation screen, followed by a sequence of images, and ends with two recall phases. **B.** In the four-image version, each image lasts for 700 milliseconds, with a 500-millisecond interval between each image. After 1500 milliseconds, the recall phase begins. To test memory performance, two of the images shown during the encoding phase are randomly selected and presented on a probe screen. The participants must identify which image appeared earlier in the four-image sequence. For each trial, memory performance is tested twice for two different pairs of images. The vertical lines are time stamps associated with ECoG recording to identify and label different phases of the task. **C.** The set of images used for the image sequence recall paradigm is also shown.

### Instantaneous Phase, Frequency, and Amplitude

We analyzed ECoG signals using bandpass filtering in conjunction with the Discrete Hilbert Transform (DHT). The raw ECoG signals were filtered in the gamma (40-150 Hz)) and theta (4-9 Hz) frequency bands using a zero-phase bandpass filter. Subsequently, we applied the DHT, which yielded an analytical function (complex function). The phase and absolute value of the analytic function from the filtered ECoG were used to calculate the instantaneous phase, frequency, and amplitude of the theta and gamma components of the ECoG signal (Madhavan, et al., 2015; Salimpour et al., 2022).

### Theta Phase Modulated Gamma Power

To quantify the gamma power modulation by the phase of the theta oscillation PAC level is estimated. To quantify PAC level, the modulation index (MI) is used (Tort et al., 2008, 2009). The estimation of MI involves filtering the signal into two components - a lower frequency band to extract the phase and a higher frequency band to extract the amplitude. The instantaneous phase of the lower frequency component and the instantaneous amplitude of the higher frequency component are then computed, using the Hilbert transform. The relationship between the phase of the lower frequency oscillation and the amplitude of the higher frequency oscillation is assessed by creating a phase-amplitude histogram, where the amplitude values are binned according to the phase at which they occur. The modulation index is then calculated from this histogram. It quantifies the degree of coupling, usually normalized to range from 0 (no coupling) to 1 (maximum coupling), by measuring the divergence of the observed amplitude-phase distribution from a uniform distribution, often using statistical measures like the Kullback-Leibler divergence. A higher modulation index indicates a stronger coupling between the phase and amplitude signals, suggesting that the phase of the low-frequency signal is coupled with the amplitude of the high-frequency signal (Tort et al., 2008, 2010). Although there is no mechanistic way to select the parameters such as the step size and the number of bins, the selections here are likely appropriate for the study hypothesis and have been used by other authors (Scheffer-Teixeira et al., 2012; Tort et al., 2009, 2010; Samiee et al., 2010; Salimpour et al., 2022).

We used this MI measure to explore the association between PAC and memory performance by analyzing the recall phase of the task. As described before (Salimpour and Anderson, 2019; Madhavan, et al., 2015), each trial consisted of two recall sections (Figure 1.B). The time of recall period starts with the presence of the probe images and ends whenever the subject presses the keypad as a response. In each recall section, the subject has a maximum of 6 seconds for pressing the key. This required us to use a variable analysis window size depending on the response time for each subject. We estimated the MI for each recall phase separately using the time window size associated with each individual subject. In order to validate the accuracy of PAC detection, a surrogate control analysis was used (see the statistical analysis session for more detail) as shown in Figure 3.B (Tort et al., 2010). To track the changes in PAC over time, we used the time-resolved phase-amplitude coupling method (Samiee et al., 2010; Salimpour et al., 2022).

**Figure 2.**
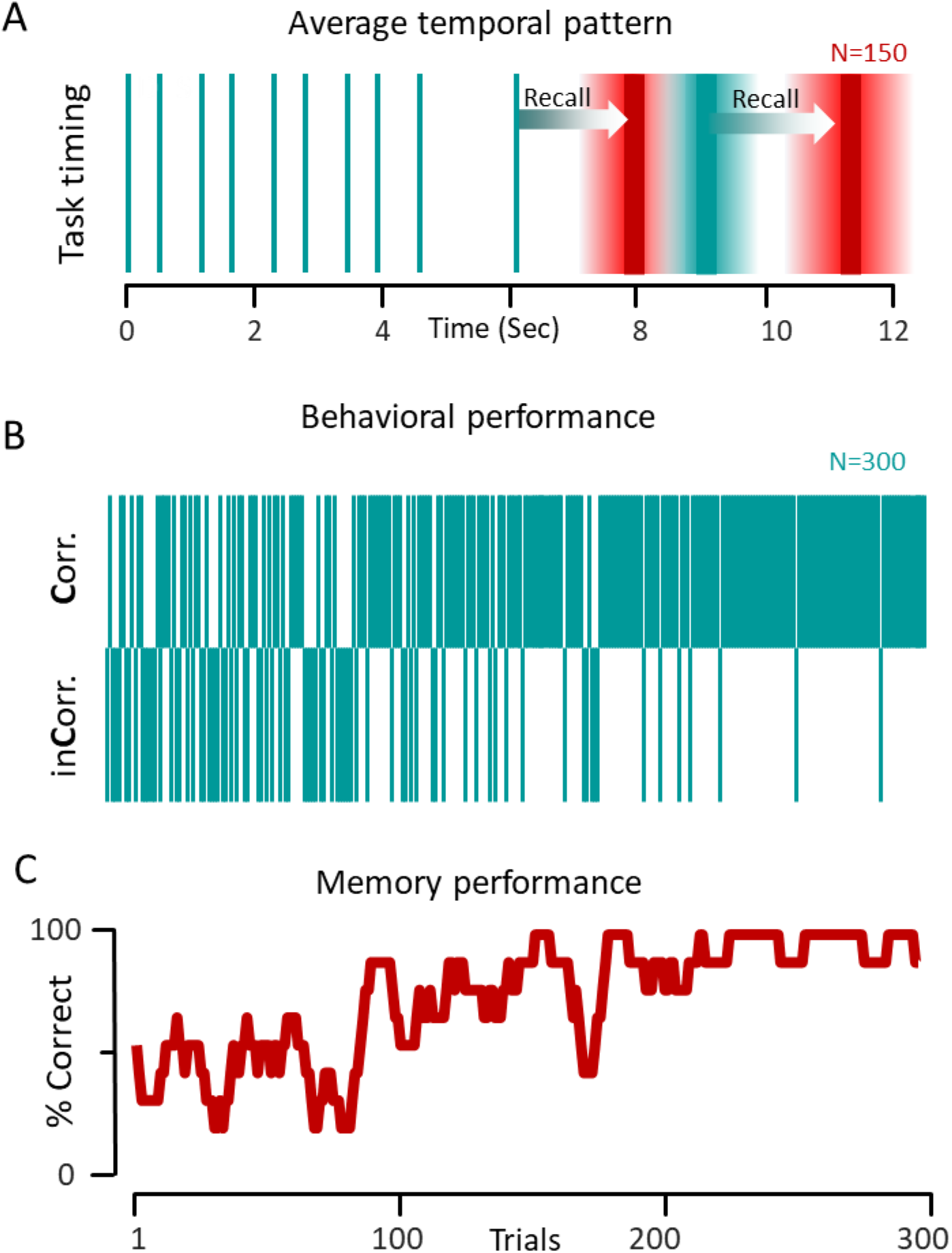
Subject’s behavioral illustration. A. The temporal pattern of choice selection for a representative subject during the behavioral task is illustrated. The green vertical lines demonstrate the time stamps for image presentation during encoding and the probe timings. The two red lines show the average reaction times, and the shaded areas are the error bars (M±Std). **B.** Behavioral profile of the representative subject showing correct and incorrect responses (n=300 responses) during the entire session (n=150 trials). **C.** Associated memory performance computed from the responses by using a sliding window of 10 responses in steps of 1 response.

**Figure 3.**
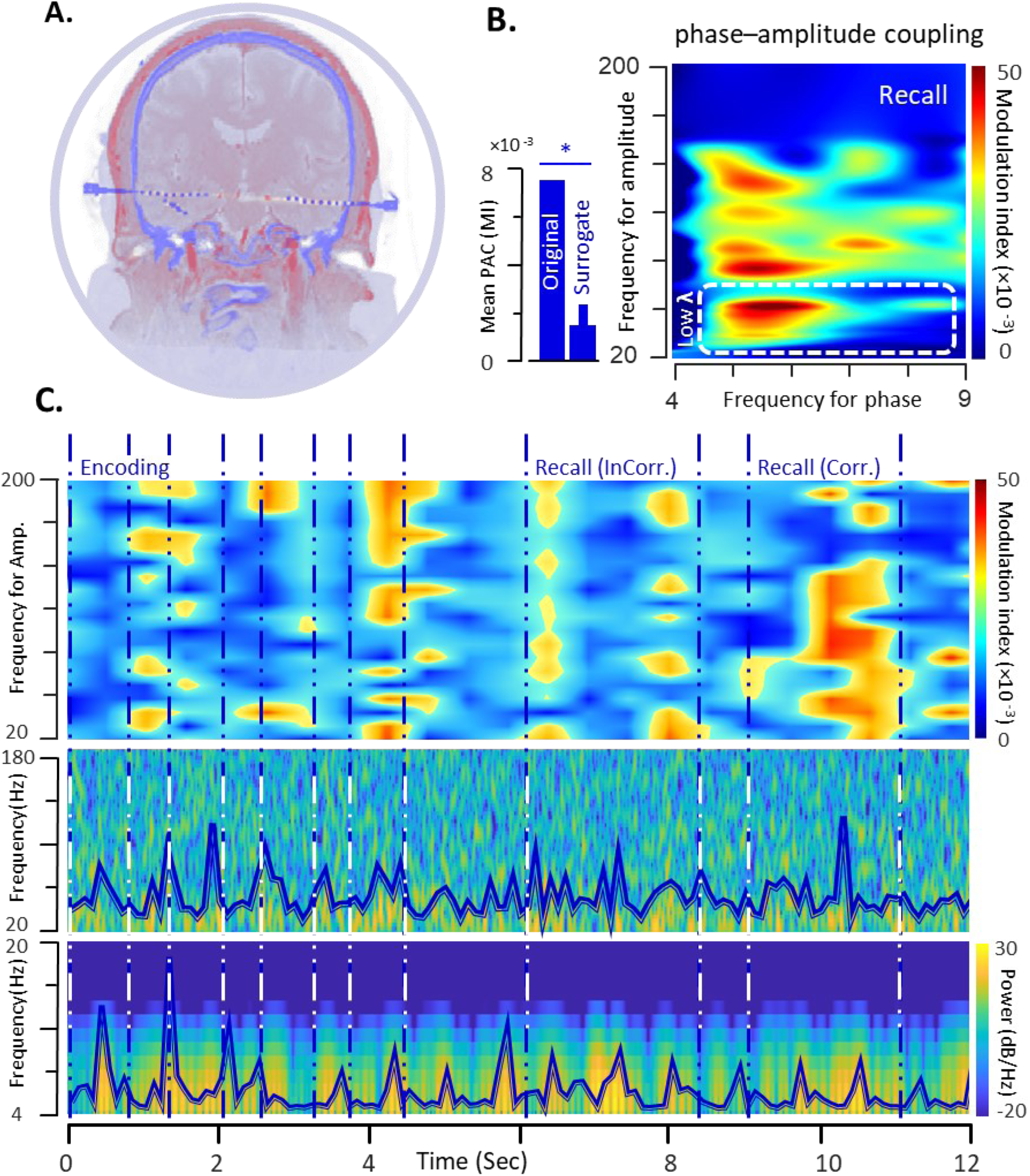
Hippocampal PAC Measurement. **A.** For the representative subject, sample electrode coverage from the left and right hippocampus was visualized using the post-op CT and the pre-op MRI. **B.** For a sample contact located in the hippocampus, the modulation index measure was used to quantify PAC in the hippocampus by analyzing the recall phase of the task. The bar chart shows the average PAC level in the recorded ECoG, and a set of randomly created (n=100) surrogate versions of the theta phase and gamma amplitude using original data. **C.** A sample trial from the representative subject with its associated dynamic PAC representation is shown. The dynamic PAC measures demonstrate the selectivity of PAC level to the behavioral response and memory performance of the subject. The associated dynamic theta and gamma range power distribution also were shown.

### Statistical Analysis

For all statistical analyses, the MATLAB Statistical package was used (R2022a). To assess the statistical significance of detected PAC changes, each experimental testing section, including a period during the retrieval phase, was analyzed using surrogate control analysis (Tort et al., 2010). We performed a statistical control analysis on each session by creating shuffled versions of the complex time series consisting of theta phase and gamma amplitude. This generated surrogate MI values, from which the MI chance distribution could be inferred (Figure 4.B). The Wilcoxon signed-rank test was employed for checking the significance of mean MI changes in paired tests, and a p-value of less than 0.05 was considered statistically significant (Tort et al., 2008, 2009, 2010). Additionally, the Pearson’s linear correlation coefficient was used as a statistical test to check whether the mean MI measured from a set of electrodes in a specific brain region had a significant correlation to the average memory performance at the 5% significance level.

**Figure 4.**
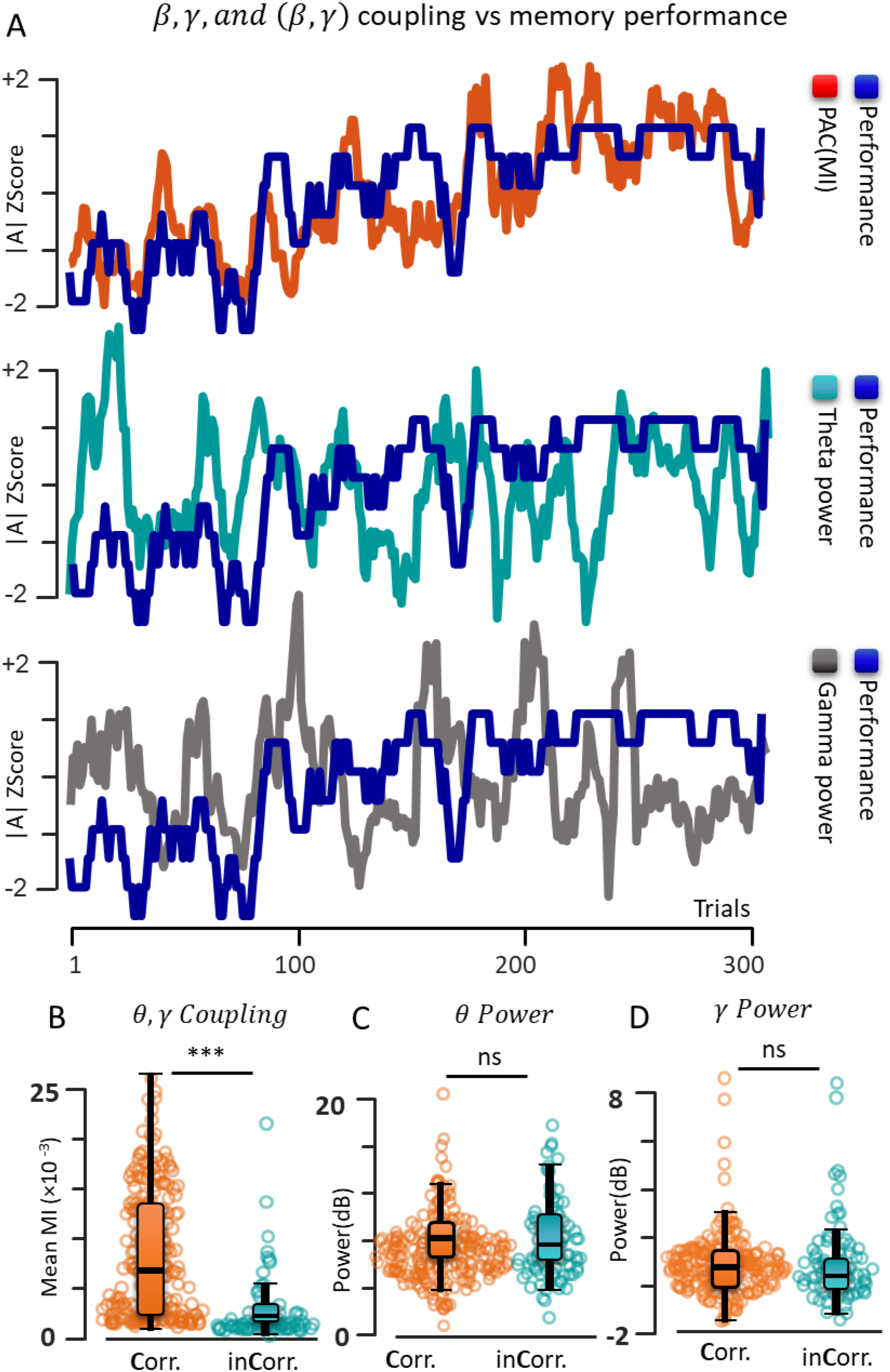
*β*,*γ*,and (*β*,*γ*) coupling vs memory performance. **A.** The temporal pattern of memory performance is compared with the associated mean modulation index, theta power, and Gamma power from hippocampus recording during n=300 trials for a representative subject. In the top panel, the behavioral profile and the pattern of PAC evolution are shown. In the middle and bottom panels similar visualization is used for behavioral profile and the pattern of changes in theta and gamma power. **B.** The level of PAC, power of theta oscillation, and gamma band power are compared for the correct and incorrect responses. The statistical analysis show significant difference between the level of PAC during correct and incorrect responses, However there are no differences between theta and gamma power during correct and incorrect responses.

## 3 Results

We analyzed ECoG activity from the intracranial grid, strip, and depth electrodes implanted in 10 epilepsy subjects while they performed an image sequence encoding and recall task (Figure 1.A) comprised of a sequence of four images selected randomly from a set of pictures including faces and buildings (Figure 1.C). We used the data from some of our previously published data sets (Salimpour and Anderson, 2019; Madhavan, et al., 2015) and added more recordings from our recently collected data. In general, the task involved encoding and recalling the temporal order of sequentially presented images. After preprocessing the recorded signals, the frequency spectrum and the phase-amplitude coupling between the amplitude of the oscillations in the gamma range (20-200 Hz) and the phase of the rhythms in the theta band (4-9 Hz) in each trial were measured. We then investigated whether CFC in the PAC in the gamma and theta frequency bands correlated to memory performance in the hippocampus.

### Memory Performance

For measuring and quantifying memory performance, we used a similar paradigm from our previously published study (Salimpour and Anderson, 2019; Madhavan, et al., 2015). In brief, a sequence of four images was presented, and a choice screen was displayed showing two randomly selected images out of the four images after a short delay. Participants were asked to remember the image sequence, and then they had to report the order in which the images had been presented by pressing the appropriate buttons on a gamepad (Figure 1). In Figure 2.A, the average temporal pattern of image presentation and choice selection for a representative subject during the behavioral task is illustrated. In Figure 2.B, the behavioral performance of a representative participant during the visual sequence memory task is shown (1 represents a correct trial and 0 codes for an incorrect response, n=300 trials) at each trial and in Figure 2.C the associated memory performance curve is calculated by using a sliding window of ten trials stepped by one trial.

### Modulation Index-based PAC Measure

We analyzed ECoG from the contacts in the deeply implanted electrodes with hippocampal coverage (Figure 3.A) and estimated the mean PAC level using the modulation index method (Figure 3). We computed the PAC level between the phase of theta (4-9 Hz) and the amplitude of the low gamma frequency range (40-150 Hz) which is mostly associated with the retrieval phase of memory-related activities in the hippocampus (Colgin et al., 2009; Colgin and Moser, 2010) (Figure 3.B). Additionally, a surrogate control analysis was used to verify the validity of PAC detection (Figure 3.B). In Figure 3.C, a sample trial from representative subject is shown. The dynamic PAC measures demonstrates the selectivity of PAC level to the behavioral response and memory performance of the subject. The associated theta and gamma range time-frequency based power distribution is also demonstrated (Figure 3.C)

### *β*,*γ*,and (*β*,*γ*) coupling vs memory response and performance

To demonstrate PAC in the form of a coupling between the amplitude of the gamma oscillation and the phase of the theta rhythm, the temporal pattern of the memory performance curve was compared with the associated modulation index waveform at a recording site in the hippocampus (Figure 4.A, for a representative subject). We explored the relationship between memory performance and the coupling between theta phase and gamma amplitude during two recall phases of the task. Each recall phase was considered as an individual trial and the percentage of the correct recall estimated using sliding windows of ten trials shifted in steps of one trial (Figure 4.A). We repeated a similar analysis to demonstrate the association between the power of the theta and gamma oscillation and the memory performance (Figure 4.A). By calculating mean MI, mean theta power, and mean gamma power during the recall period, we conducted a statistical analysis to determine the significance of changes in any of these measures during correct and incorrect responses. In Figure 5.B, the trial-by-trial estimation of mean MI, mean theta power, and mean gamma power during the recall period associated with correct and incorrect responses. The statistical analysis shows a significant change in the level of coupling between the phase of the theta and the amplitude of the gamma, but no significant changes detected in theta and gamma power individually.

**Figure 5.**
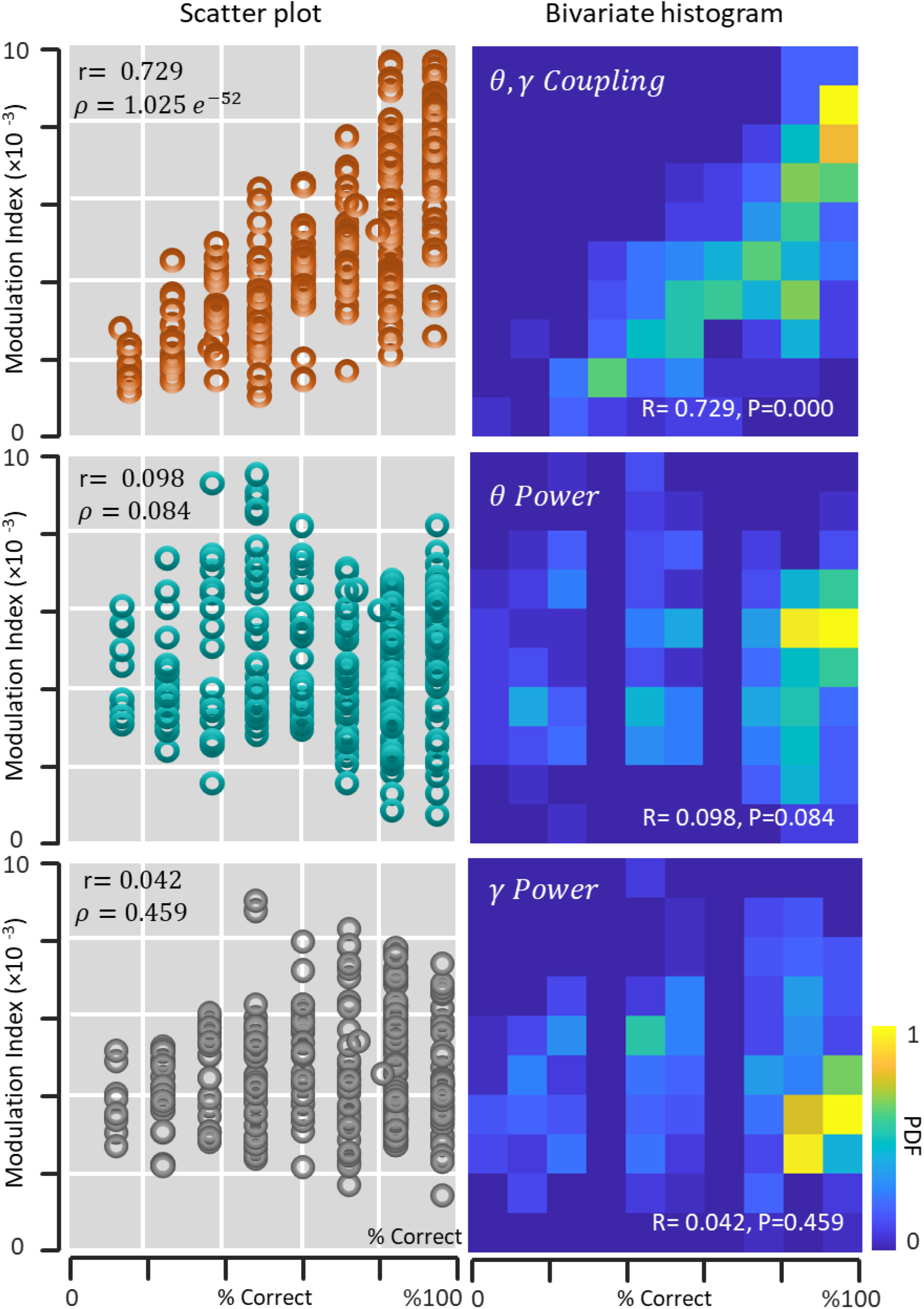
Linear correlation between *β*,*γ*,and (*β*,*γ*) coupling, and task performance: To quantify and measure the linear correlation between modulation index, theta power, gamma power, and memory performance, the scatter plot of each measure over each mean performance percentage is used. Additionally, The bivariate histogram is used to visualize the distribution of memory performance and modulation index, theta power, and gamma power. Pearson’s correlation coefficient quantifies correlations between variables. The correlation between the average MI value (top panel), mean theta power (middle panel), and mean gamma power (bottom panel) over each mean performance percentage are shown.

The trial-by-trial analysis of enabled us to quantify and measure the linear correlation between modulation index, theta power, gamma power, and memory performance. The scatter pattern of each measure over each mean performance percentage is used to investigated the possible correlation (Figure 5). Additionally, The bivariate histogram is used to visualize the distribution of memory performance and modulation index, theta power, and gamma power. Pearson’s correlation coefficient quantifies correlations between variables. There is a significant correlation between the mean performance percentage and average MI value. However, we did not detect any correlation between memory performance and mean theta power or mean gamma power (Figure 5).

### Group Temporal and Behavioral Performance

In Figure 6, all derived subject behavioral information and associated PAC measures are summarized. ECoG segments for estimating PAC (in MI measures) between the amplitude of gamma and the phase of theta are extracted from electrode contacts by recording from the time of probe presentation until the response was recorded by the gamepad, which here is called the recall phase of the task. Figure 6.A shows the average time windows with associated variabilities used here for the detection and estimation of PAC. The associated memory performance was computed for all subjects using a sliding window of 10 trials in steps of one trial (the green curve shows the memory performance of the representative subject).

**Figure 6.**
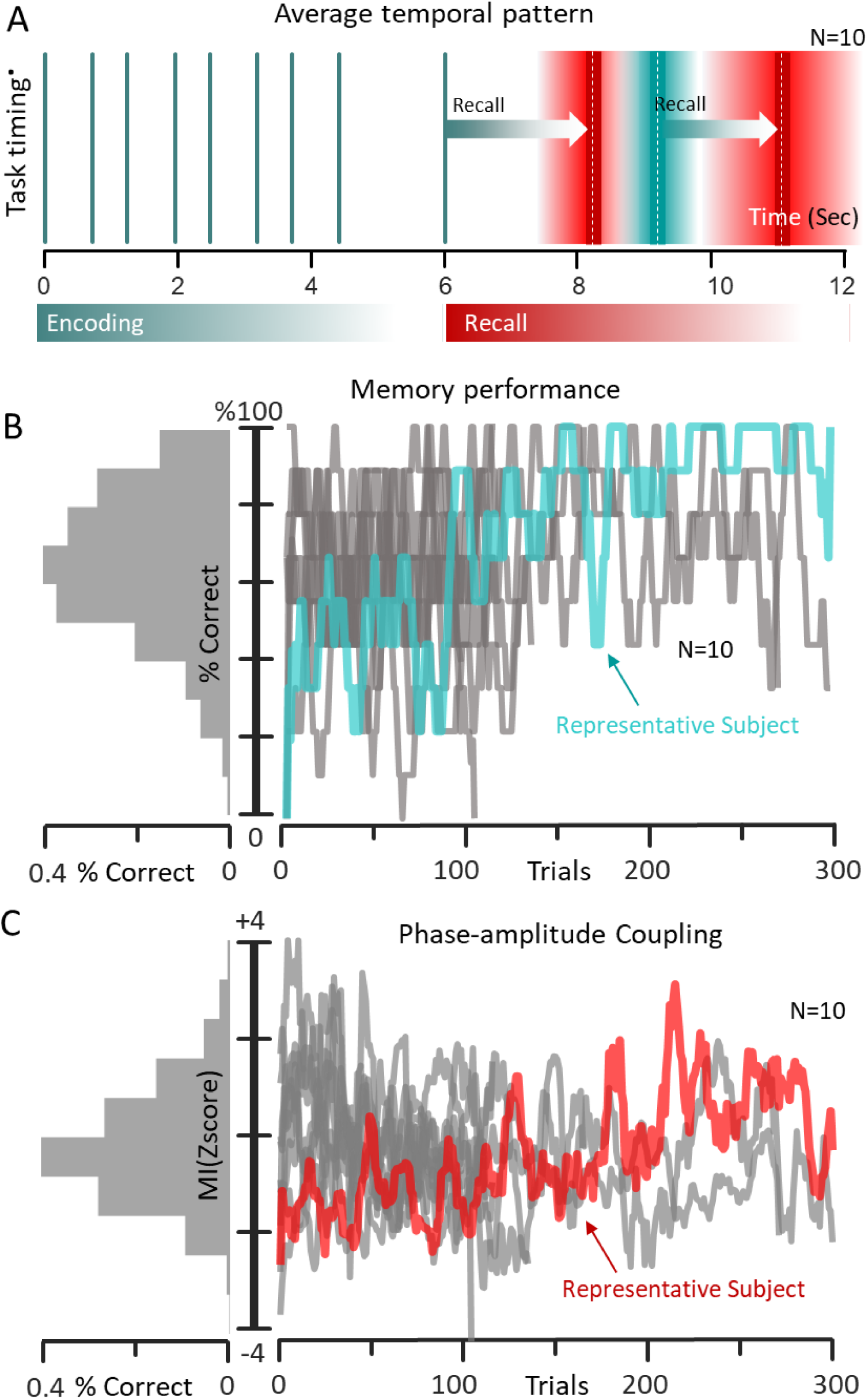
Memory performance of the subjects. **A.** Temporal patterns of the stimulus presentation and choice selection from all participants during memory testing are illustrated. Similar to the representative subject data, the green vertical lines demonstrate the image presentation times during the encoding phase of the task (same values for all subjects) and the probe timing of the two consecutive recalls. The two red lines demonstrate the average reaction times across all subjects, and the shaded areas demonstrate the standard error of the mean values. The reaction times for each trial are used as the time windows for estimating the mean modulation index, mean theta power, and mean gamma power during the recall phase of the task. **B.** Associated memory performance was computed for all subjects using a sliding window of 10 trials in steps of one trial (the red curve shows the memory performance of the representative subject). The histogram of the mean performance in each percentile for all subjects is demonstrated on the left side of the figure. **C.** Similar to the memory performance (panel B), the mean MI level was computed for all subjects using a sliding window of 10 trials in steps of one trial (the red curve shows the mean MI level of the representative subject). The histogram of the mean MI level in each performance percentage for all subjects is demonstrated on the left side of the figure.

Figure 6.B summarizes the behavioral data and the histogram of the mean performance in each percentile for all subjects is demonstrated on the left side of the figure as a marginal distribution. Similar to the memory performance in Figure 6.B, the mean MI levels were computed for all subjects using a sliding window of 10 trials in steps of one trial (the red curve shows the mean MI level of the representative subject) and summarized in Figure 6.C. The marginal distribution of the mean MI level in each performance percentage for all subjects histogram is demonstrated on the left side of the figure.

### Aggregate Electrode Positions on the Hippocampus

In this study, we were interested in memory-related structures, but our focus was on the contacts located in the hippocampus. Precise anatomical locations of the electrodes were estimated using the pre-implant MRI and a post-implant CT scan, and finally, the explained procedure in the method session was used for anatomical labeling of the regions of interest (Fan et al., 2016; Griffin et al., 2018). In Figure 7.A, the aggregate electrode positions from all implanted electrodes located inside the hippocampus across the patients are rendered on a normalized hippocampus (Griffin et al., 2018).

**Figure 7.**
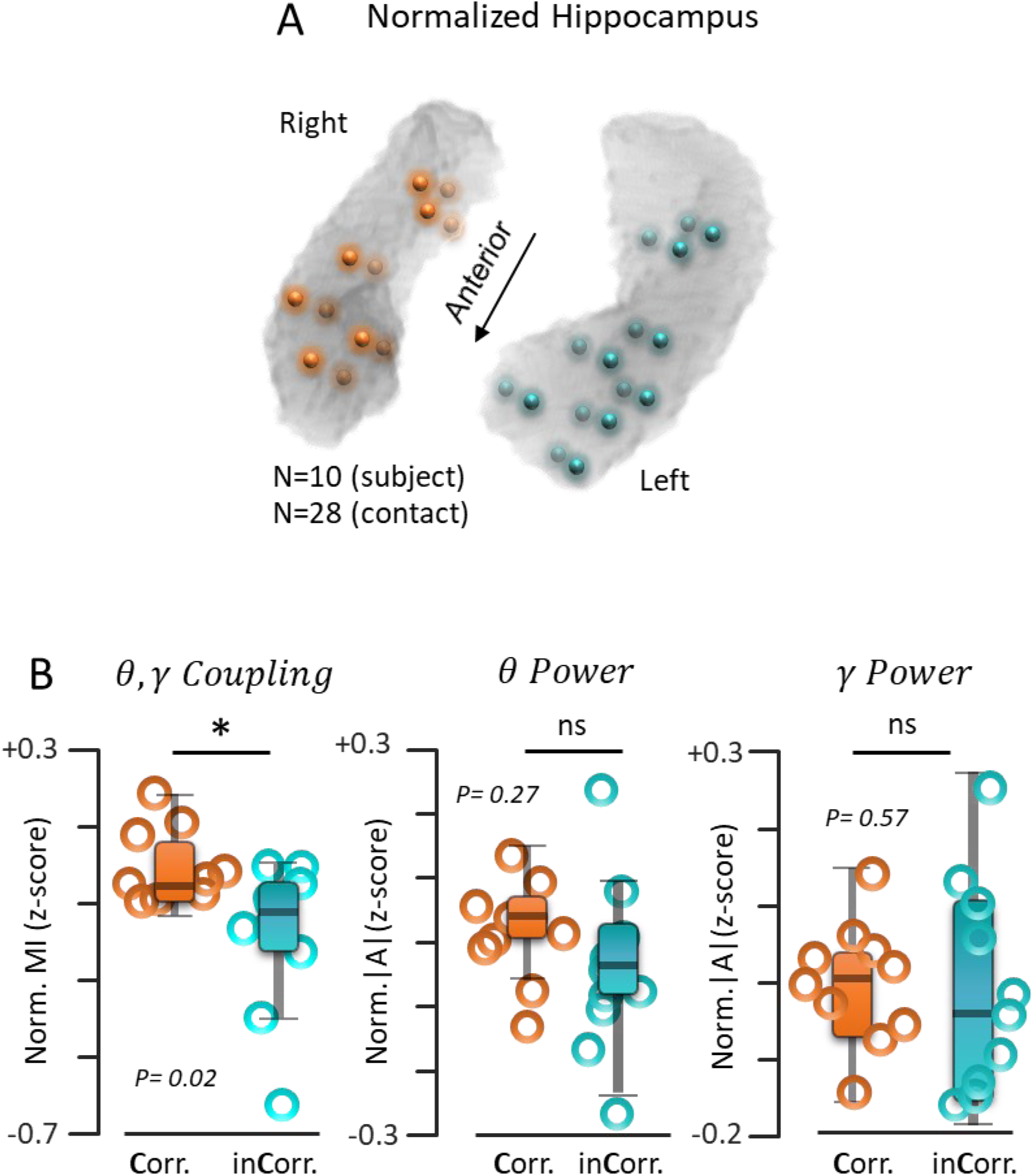
Hippocampal activity and memory responses. A. Accumulative locations of all contacts across the subjects used in the hippocampal theta power, gamma power, and theta-gamma phase- amplitude coupling analysis. **B.** The correct and incorrect responses extracted from memory performance were used to test the significance of changes in theta power, gamma power, and theta- gamma phase-amplitude coupling level associated with each response. Only the PAC level significantly changes between correct and incorrect responses.

### Hippocampal activity and memory responses

In Figure 7.B, we summarized the results of analyzing the memory responses of our subjects (N=10) and related hippocampal activities (N=32 contacts located inside the hippocampus). For each patient, we averaged theta power, gamma power, and mean MI level estimated from hippocampal activities associated with memory responses. The correct and incorrect responses extracted from memory performance were used to test the significance of changes in theta power, gamma power, and theta-gamma phase-amplitude coupling level associated with each response. Only the PAC level significantly changes between correct and incorrect responses.

### Hippocampal activity and memory performance

In order to investigate the linear correlation between modulation index, theta power, gamma power, and memory performance in group level, we extended the similar analysis we did for the representative subject (Figure 5). In Figure 8, we summarized the hippocampal activity dependence on memory performance using the average modulation index measures, theta power, and gamma power with respect to the normalized memory performance across all participants. To illustrate the hippocampal response at the group level and in the form of a mean modulation index with respect to the memory performance, the mean of the normalized MI (z-score) for all selected contacts across all subjects is averaged in each percentage of the mean memory performance. In Figure 8.A, the graph represents the relationship between memory performance and modulation index. The horizontal axis shows the average behavior of all subjects at each percentage of memory performance, while the vertical axis shows the average of the mean of normalized MI (z-score) values for all contacts associated with each percentage of memory performance. Additionally, the marginal histograms demonstrate the distribution of the data for each variable individually. The bivariate histogram is a useful tool for visualizing the distribution of memory performance and hippocampal activity at the group level. Pearson’s correlation coefficient indicates strong correlations between the mean of normalized MI and memory performance (Figure 8.B) . However, the bivariate histogram shows that there is no correlation between memory performance and theta power (Figure 8.C), as well as no correlation between memory performance and gamma power (Figure 8.D).

**Figure 8.**
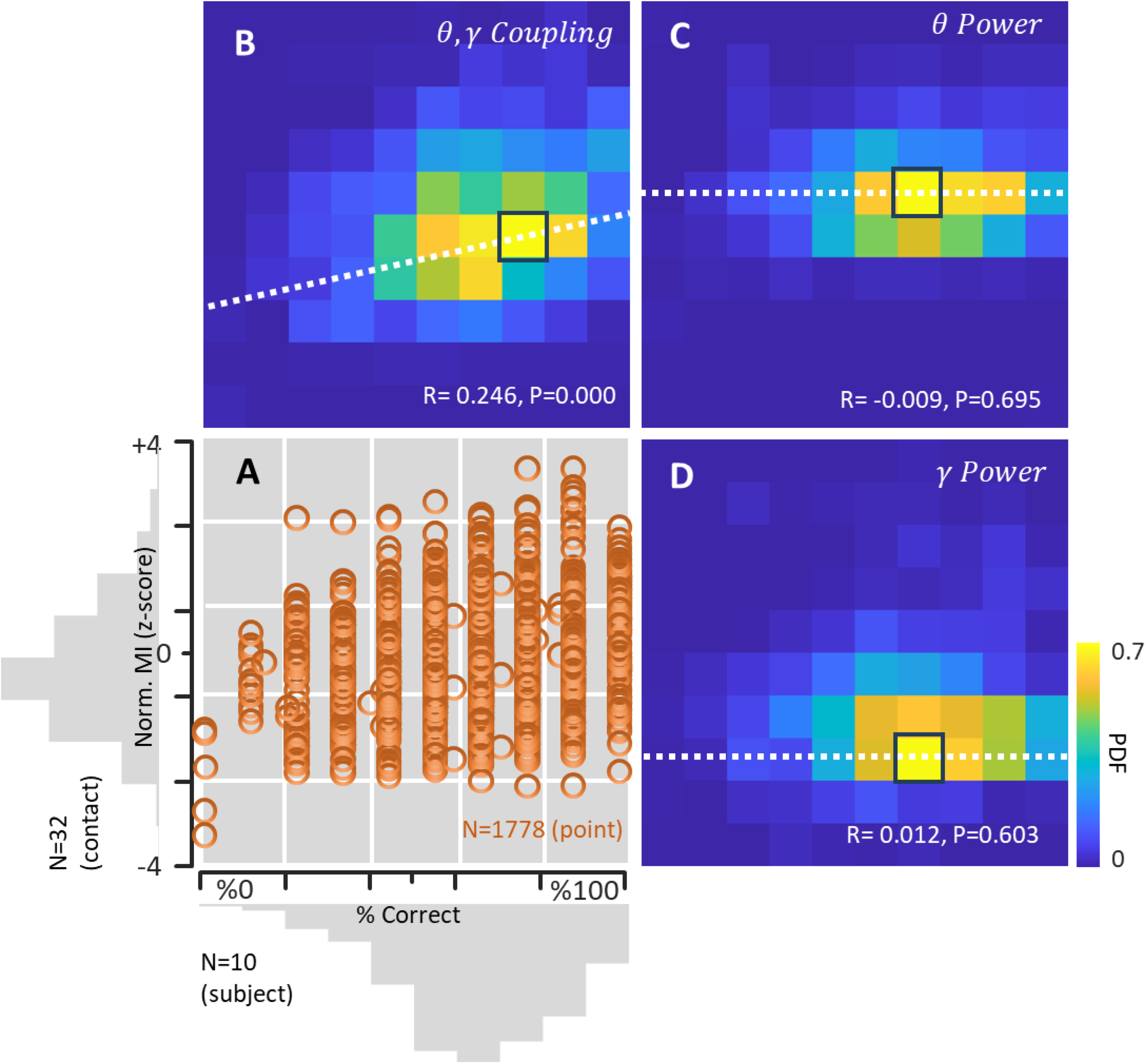
Hippocampal activity and memory performance. Hippocampal activity dependence on memory performance is demonstrated with the average modulation index measures, theta power, and gamma power with respect to the normalized memory performance. **A.** To illustrate the hippocampal response at the group level and in the form of a mean modulation index with respect to the memory performance, the mean of the normalized MI (z-score) for all selected contacts across all subjects is averaged in each percentage of the mean memory performance. In this representation, the horizontal coordinates are the average of all subjects’ behavior in each percentage of memory performance. The vertical coordinates are the average of the mean of normalized MI (z-score) values for all contacts associated with each percentage of the memory performance. The main plot shows the relationship between the memory performance and modulation index and the marginal histograms show the distribution of the data for each variable. **B.** The bivariate histogram is used to visualize the distribution of memory performance and hippocampal activity in the group level. Pearson’s correlation coefficient shows strong correlations between two variables. **C.** The bivariate histogram shows no correlation between memory performance and theta power. **D.** The bivariate histogram shows no correlation between memory performance and gamma power.

## 4 Discussion

In this research effort, we investigated the functional roles of theta rhythm, gamma oscillation, and their cross-coupling in the human hippocampus during working memory tasks. The results from our study prove a stronger functional role for the theta-gamma cross-coupling in comparison to the power of theta and gamma individually in the human hippocampus during working memory performance. Specifically, the level of coupling between theta-phase and gamma amplitude is correlated with the accuracy of memory performance. These results support the hypothesis that the theta–gamma cross-frequency coupling in the form of PAC contributes to the functional neurobiology of the memory process in the human hippocampus (Lisman and Idiart 1995; Canolty et al. 2006; Axmacher et al. 2010; Fuentemilla et al. 2010; Jacobs and Kahana 2009; Mormann et al. 2005; Sauseng et al. 2009; Tort et al. 2009; Tort et al. 2008).

We decided to use a cross-frequency coupling calculation as the neurophysiology metric rather than simple rhythmic activities to correlate with memory performance for the following reasons: Theta rhythms in the hippocampus are not limited to memory-related activities and might emerge during other sensory and cognitive processes such as sensory information processing, motor activity, and rapid eye movements (Bland and Oddie 2001; Buzsaki 1996; Montgomery et al. 2008). On the other hand, gamma oscillations were found to be induced in the hippocampus under different neuronal mechanisms, mostly associated with the cell types involved in generating the oscillations including interneuronal network gamma activity, pyramidal interneuron network gamma, and sparse and persistent gamma activity which has been observed in a variety of brain activities and functions (Whittington et al. 2011). This is in agreement with our results since we did not detect a correlation between working memory and the power of theta and gamma individually.

Additionally, CFC has the potential to aid in understanding the neurobiology of working memory facilitated by the hippocampus neurophysiological activity measurements. CFC metrics, especially in the form of PAC, offer a potentially efficient tool for investigating the neuronal mechanisms underlying multi-item representation in the hippocampus. Our understanding of the neurobiology of PAC infers its origin as a temporally clustering mechanism of spikes associated with distinct theta oscillation phases which produce gamma band activity coupled to a specific theta phase (Jensen, 2001; Akam and Kullmann, 2010).

Based on hippocampal electrophysiological recordings, gamma rhythms are generally temporally nested within the theta rhythm leading to the existence of complex interactions between gamma and theta, such as CFC (Bragin et al. 1995; Montgomery et al. 2008). CFC arises in different anatomical pathways and also during memory-related behavioral tasks (Goutagny et al. 2009; Canolty et al., 2010; Buzsaki and Wang 2012; Lisman and Jensen, 2013). Most of these studies used PAC (with the modulation index measure) as a representation of CFC to quantify the complex interaction between theta and gamma rhythms and have demonstrated memory function associations with PAC in the human hippocampus (Axmacher et al., 2010; Fell and Axmacher, 2011; Lega et al. 2015; Kendrick et al. 2011; Canolty and Knight 2010).

PAC measures in the hippocampus might represent the coordination between neuronal oscillations that arise from the functional interactions between gamma and theta rhythm generators. More specifically, PAC might be used in the hippocampus to shape the temporal pattern of spiking activities in both the pyramidal cells and interneurons. This is in line with our study results since we observed a stronger functional role for the theta-gamma cross-coupling in the human hippocampus during memory-related activities.

To investigate the function of PAC in a memory-related role in the hippocampus, we used our previously published memory task and measured both ECoG and memory performance in human subjects (Salimpour and Anderson, 2019; Madhavan, et al., 2015). We found that CFC in the form of PAC between theta phase and low gamma amplitude was more pronounced and significantly increased during the probe phase of the task as compared to the theta and gamma power individually. To reduce the computational load for estimating the modulation index, we simplified the PAC detection algorithm and restricted it to computing only the phase of theta frequency (in the range 4-9 Hz) and amplitude of gamma band (in the range 40-150 Hz) activity. These findings demonstrate that cross-frequency coupling in select temporal lobe structures, including the hippocampus, correlates with memory function.

For the following reasons, our estimation of the average power of the gamma-band oscillation coupled to the phase of the theta rhythms might not be the most optimal technique. First, we are using very limited recordings of ECoG from responsive contacts. A longer analysis window would provide better cross-frequency coupling estimation. Here, we always kept the analysis window above a specific threshold. Second, we limited the frequency range for phase estimation to the theta band. This assumes that the cross-frequency coupling is produced mostly in the theta-gamma range. If the coupling happens in other frequency ranges, it would not be included in the estimation process. Finally, the cross-frequency coupling is only measured in the phase-amplitude format, and other forms of coupling may be missed in the coupling detection process (Scheffer-Teixeira et al., 2012; Tort et al., 2008, 2009).

Here, we have used a visual memory paradigm with a consecutive presentation of pictures and measured the average power of the gamma oscillation coupled to the phase of the theta rhythm in the form of an average modulation index and correlated it to subject memory performance. Our results demonstrate that the average modulation index between the phase of the theta waveform and the amplitude of gamma power is significantly correlated with memory performance and can potentially be used as a predictor of behavioral performance in similar memory tasks. Our modified MI detection algorithm is helpful for near real-time memory performance monitoring applications with potential capabilities for quantifying the later effects of planned surgical treatments, including selective laser ablation and open resections on memory structures.

## Author contributions

W.S.A. and Y.S. performed the experiments, analyzed the electrophysiological and behavioral data and wrote the manuscript. W.S.A. performed the electrode implantations and necessary neurosurgical procedures. Y.S. ran the experiments and recorded both the electrophysiological and behavioral data; both authors edited the manuscript.

